# Generation of *Drosophila* attP containing cell lines using CRISPR-Cas9

**DOI:** 10.1101/2021.03.19.436223

**Authors:** Daniel Mariyappa, Arthur Luhur, Danielle Overton, Andrew C. Zelhof

## Abstract

The generation of *Drosophila* stable cell lines have become invaluable for complementing *in vivo* experiments and as tools for genetic screens. Recent advances utilizing attP/PhiC31 integrase system has permitted the creation of *Drosophila* cells in which recombination mediated cassette exchange (RMCE) can be utilized to generate stably integrated transgenic cell lines that contain a single copy of the transgene at the desired locus. Current techniques, besides being laborious and introducing extraneous elements, are limited to a handful of cell lines of embryonic origin. Nonetheless, with well over 100 *Drosophila* cell lines available, including an ever-increasing number CRISPR/Cas9 modified cell lines, a more universal methodology is needed to generate a stably integrated transgenic line from any one of the available *Drosophila melanogaster* cell lines. Here we describe a toolkit and procedure that combines CRISPR/Cas9 and the PhiC31 integrase system. We have generated and isolated single cell clones containing an Actin5C::dsRed cassette flanked by attP sites into the genome of Kc167 and S2R+ cell lines that mimic the *in vivo* attP sites located at 25C6 and 99F8 of the *Drosophila* genome. Furthermore, we tested the functionality of the attP docking sites utilizing two independent GFP expressing constructs flanked by attB sites that permit RMCE and therefore the insertion of any DNA of interest. Lastly, to demonstrate the universality of our methodology and existing constructs, we have successfully integrated the Actin5C::dsRed cassette flanked by attP sites into two different CNS cell lines, ML-DmBG2-c2 and ML-DmBG3-c2. Overall, the reagents and methodology reported here permit the efficient generation of stable transgenic cassettes with minimal change in the cellular genomes in existing *D. melanogaster* cell lines.

## Introduction

Transgenesis is a powerful and proven technique in *Drosophila* biology, both *in vivo* and *in vitro* tissue culture cells. Over time, the techniques utilized have been modified and refined to increase the efficiency of generating stable transgenic lines (Groth *et al*. 2004; Venken *et al*. 2006; Bischof *et al*. 2007; Venken and Bellen 2007; Pfeiffer *et al*. 2010). With respect to *Drosophila* cell lines, the initial generation of stably integrated transgenes has previously relied on the transfection of a plasmid of interest in combination with a selectable marker (neomycin or hygromycin) as a dual plasmid system or with dual expression cassettes in a single vector or using a multicistronic vector (Bourouis and Jarry 1983; Koelle *et al*. 1991; Iwaki *et al*. 2003; Iwaki andCastellino 2008; Gonzalez *et al*. 2011). Alternatively P-element mediated transformation has also been attempted (Segal *et al*. 1996). The downsides to these approaches have included random genomic insertions, generation of unstable tandem arrays (Cherbas *et al*. 1994), possibly skewed selection of drug resistance over transgenic expression (Gonzalez *et al*. 2011) and ultimately leading to inconsistent expression of the introduced transgenes (Spradling and Rubin 1983).

With the advent of phiC31 site directed insertion and recombination mediated cassette exchange (RMCE) in *Drosophila* (Groth *et al*. 2004), the generation of single copy transgenes into a single genomic locus was possible in cells (Neumuller *et al*. 2012) and subsequently two different methodologies have been developed (Cherbas *et al*. 2015; Manivannan *et al*. 2015). The first strategy was to derive cell lines from a *Drosophila melanogaster* stock containing a P-element carrying a mini-white flanked by attP sites that allowed RMCE combined with ubiquitously expressed Ras^V12^ to aid in the recovery of a stable cell line (Manivannan *et al*. 2015). Two embryo-derived stable cell lines, Ras-attP-L1 and Ras-attP-L2, were generated using this strategy. Crucially, subsequent transfection of a RMCE competent exchange cassette with phiC31 integrase led to creation of cell lines containing a single copy of the transgene (Manivannan *et al*. 2015). The second approach utilized P-element transposition mediated insertion of a cassette with Act5C::GFP flanked by attP, directly into Kc167 and Sg4 cells (Cherbas *et al*. 2015). Transformants were selected and cell lines containing single P-element insertions were generated. As with the Ras-attP lines, RMCE resulted in creation of several cell lines (Sg4-PP and Kc167-PP) with a single copy of the transgene.

Both the approaches described above have their advantages. With the Ras-attP lines, constitutive GAL4 expression can be leveraged for transgenic expression and the genomes of the Ras-attP lines are not as complex as the other established lines (Lee *et al*. 2014; Manivannan *et al*. 2015). The use of ModENCODE lines to derive the Kc167-PP and Sg4-PP lines and the availability of a multitude of insertion sites are notable advantages (Cherbas *et al*. 2015). Nevertheless, with either approach expanding the technology to existing or novel cell lines harboring attP sites at identical loci is extremely challenging in terms of resources and time.

Subsequently, CRISPR-Cas9 based approaches have revolutionized genome editing (Bier *et al*. 2018). Utilizing the CRISPR-Cas9 genome editing strategy, we have created cell lines with RMCE compatible attP sites at loci in sync with *in vivo* attP sites utilized to generate transgenic flies. The tools created here can potentially be used to generate attP sites in novel or existing cell lines at the same genetic loci with minimal change in the cellular genome. We present data outlining the creation of a S2R+ line with an attP cassette at 99F8 on the 3^rd^ chromosome and the successful RMCE with an associated attB plasmid tool kit we have developed. Another genomic locus, 25C6 on the 2^nd^ chromosome was similarly modified and tested for RMCE. In addition, we demonstrate that, attP insertions at both 25C6 and/or 99F8 loci can be created in other cell lines, specifically in embryonic (Kc167) and CNS (ML-DmBG2-c2 and ML-DmBG3-c2) lines. Eventually, the attP insertion cassette will be expanded to include more of the existing *Drosophila* cell line collection. Furthermore, all the reagents generated are available to the *Drosophila* research community to either independently create attP containing cell lines or utilize the created attP lines reported here for RMCE to insert the transgene of interest.

## Materials and Methods

### Plasmid construction and molecular biology

The loci chosen for insertion of attP sites was arrived upon by assessing the Rainbow Transgenics information (https://www.rainbowgene.com/rtf-line-collection/) on *in vivo* attP sites demonstrated to produce transformants and having high viability. Upon finalizing the target loci, guide RNA (gRNA) candidates were determined using the tool at http://tools.flycrispr.molbio.wisc.edu/targetFinder/. For the site at 99F8, a single gRNA was cloned into the pU6-3-gRNA vector (Drosophila Genomics Resource Center, DGRC Cat # 1362) as described (Gratz *et al*. 2013). For the site at 25C6 locus, two gRNAs were used and cloned independently into the pU6-3-gRNA vector. Briefly, 25C6 (2L: 2L:5108448..5108448) and 99F8 loci targeting (3R:30553233..30553255) sense and antisense oligonucleotides (Table 1) with BbsI overhangs were annealed and ligated with gel-purified, BbsI digested, dephoshorylated pU6-3-gRNA vector. A single clone from the ligation was selected on sequence confirmation of the presence of the gRNA insert.

**Table 1.**
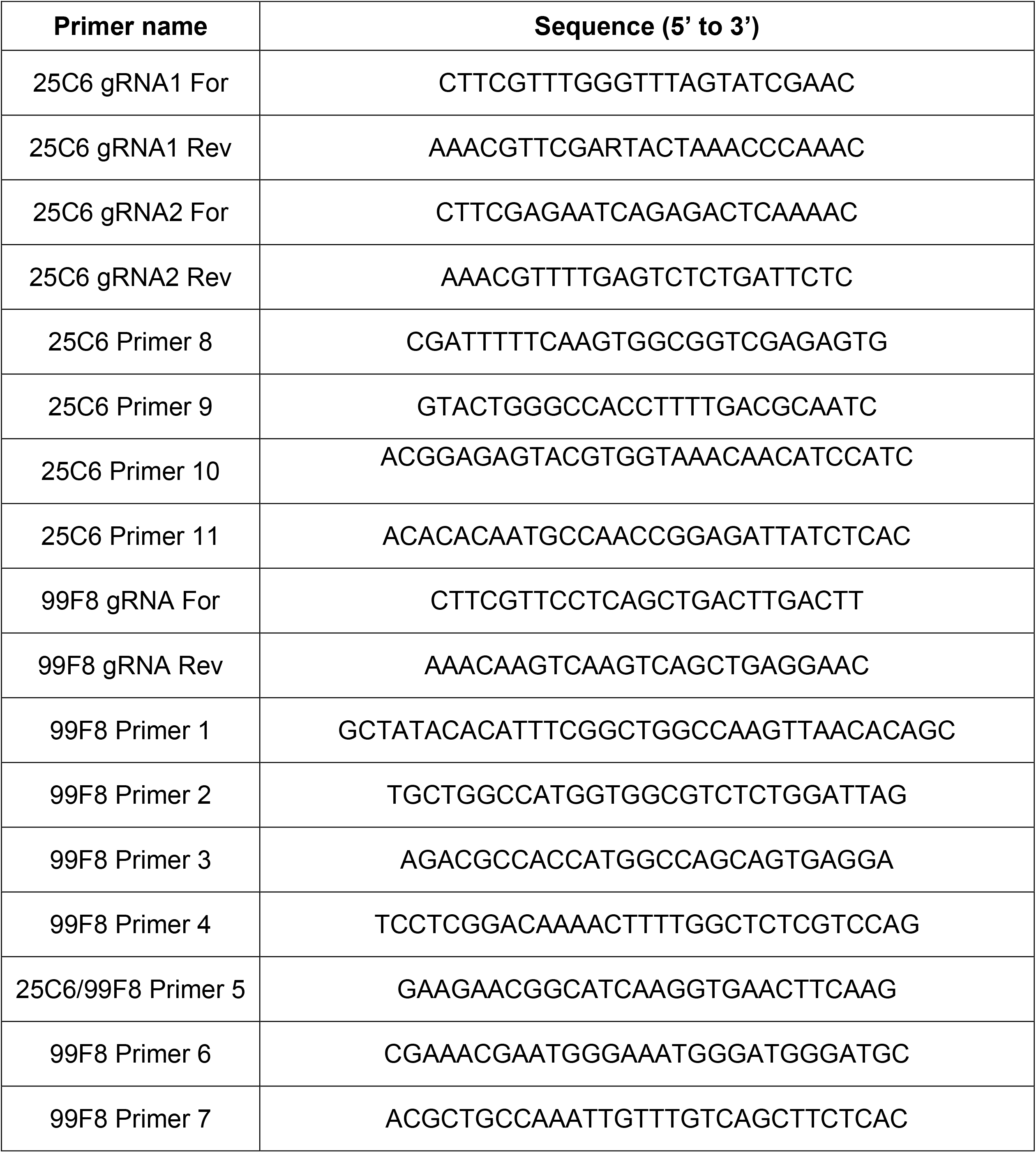
Primers used in the study.

The donor template for the 25C6 exchange cassette, pUC57-25C6-5′HR-attP>>Act5C::dsRed<<attP-3′HR (DGRC Cat# 1543) was custom synthesized by Synbio Technologies Inc., NJ. Donor template for the 99F8 exchange cassette was assembled into a pBluescript (pBS, Stratagene) vector backbone using Gibson Assembly to create pBS-99F8-5’HR-attP>>Act5C::GFP<<attP-3’HR (DGRC Cat# 1544) vector. The replacement donor was cloned into BsgI digested pBS with the 5’ and 3’ homology arms, which were 0.96 kb and 1.14 kb respectively, flanking the attP>>Act::dsRed<<attP insert amplified from pUC57-25C6-5′HR-attP>>Act5C::dsRed<<attP-3′HR. Sequencing confirmed the correct assembly of pBS-99F8-5’HR-attP>>Act5C::GFP<<attP-3’HR vector. The vectors for recombination mediated cassette exchange (RMCE), pUC57-attB>>MCS-Act5C::EGFP<<attB (DGRC Cat# 1545) and pUC57-attB>>MCS-MT::EGFP<<attB (DGRC Cat# 1546) were custom gene synthesized by Synbio Technologies Inc., NJ.

### Cell lines and transfections

Detailed cell culture protocols have been described previously (Luhur *et al*. 2019a; Luhur *et al*. 2019b). Briefly, *Drosophila* S2R+ (CVCL_Z831, DGRC Cat # 150), S2 DGRC (CVCL_TZ72, DGRC Cat # 6) cells were grown in M3 media (Sigma) containing Bactopetone, yeast extract and supplemented with 10% fetal bovine serum (M3+BPYE+10%FBS). Embryonic Kc167 (CVCL_Z834, DGRC Cat #1) cells were grown in CCM3 (GE Life Sciences) medium. Larval central nervous system derived cell lines ML-DmBG2-c2 (CVCL_Z719, DGRC Cat #53) and ML-DmBG3-c2 (CVCL_Z728, DGRC Cat #68) were grown in M3+BPYE+10%FBS supplemented with 10 µg/ml insulin.

For the CRISPR-Cas9 transfections, 2 ml (1.5× 10^6^/ml) cells plated in a well of a 6-well dish were transfected with 300 ng of Act::Cas9 (Addgene #62209) and the respective gRNA and repair templates for the 25C6 (75 ng pU6-3-25C6-gRNA1 (DGRC Cat# 1547), 75 ng pU6-3-25C6-gRNA2 (DGRC Cat# 1548) and 150 ng pUC57-25C6-5′HR-attP>>Act5C::dsRed<<attP-3′HR) and 99F8 (150 ng pU6-3-99F8-gRNA (DGRC Cat# 1549) and 150 ng pBS-99F8-5’HR-attP>>Act5C::GFP<<attP-3’HR) loci using Effectene Transfection reagent (QIAGEN) as per the manufacturer’s recommendation. Transfected cells were cultured for 10-20 days to amplify the cultures to two 100 mm dishes before bulk or single cell sorting the cells.

For RMCE transfections, 2 ml (1.5× 10^6^/ml) cells plated in a well of a 6-well dish were transfected with 300 ng of either pUC57 attB>>MCS-Act5C::EGFP<<attB or pUC57 attB>>MCS-MT::EGFP<<attB and 200 ng Act::phiC31 integrase (DGRC Cat # 1368), using Effectene Transfection reagent (QIAGEN) as per the manufacturer’s recommendation. Transfected cells were cultured for at 10-20 days to amplify the cultures to two 10 cm dishes before bulk sorting the cells.

### Fluorescence activated cell sorting

Cells were bulk- or single-cell sorted into wells of 24-well or 96-well plates, respectively. The S2 lines were sorted into wells containing S2 conditioned medium prepared as described previously (Housden *et al*. 2015). All the other cell lines were sorted into the respective standard media containing 1.5× 10^6^/ml irradiated, non-transfected feeder cells of similar origin (Cherbas *et al*. 2015). After 10-20 days post-transfection, cells were sorted using fluorescent-activated cell sorting (FACS) on a FACS Aria II (IUB Flow Cytometry Core Facility). Live cells were sorted by exclusion of DAPI, detected by excitation with a 407-nm laser. To detect EGFP or dsRed positivity, a 488-nm or a 561-nm laser, was used respectively. In the case of S2R+ and Kc167 lines, the EGFP positive bulk sorted cells were amplified to populate two 10 cm dishes before subjecting to single-cell sorting.

### Genomic PCR and single cell sorting

After bulk sorting, when the sorted cells reached to 60-75% confluency, the cultures were serially expanded to a 10 cm dish. During the earliest transition, a small fraction of the cells was harvested for genomic DNA (gDNA) isolation. The harvested cells were centrifuged and washed once with phosphate buffered saline. Genomic DNA was extracted from the phosphate buffered saline (PBS) washed pellet using the Zymo Quick-DNA^™^ MiniprepPlusKit (D4068/4069). PCR reactions were performed to diagnose and confirm CRISPR-Cas9 mediated homologous recombination (Primers pairs 1+2, 3+4 or 1+4, Figure 2, Table 1) or RMCE (Primer pairs 5+6 or 5+7, Figure 3, Table 1). Sequences of all the primers used in this study are listed in Table 1. All the potentially positive amplicons of the expected molecular weights were sequence confirmed. The single-cell clones obtained after FACS sorting were observed for growth, visually checked for single-cell clonality using a Nikon Eclipse TS100 brightfield microscope and dsRed expression using a Zeiss AX10 microscope. When the sorted, dsRed positive single-cell colonies reached to 60-75% confluency, they were passaged onto a well of a 24-well dish, concomitantly harvesting gDNA and performing PCRs to diagnose and confirm CRISPR-Cas9 mediated homologous recombination (Primers pairs 1+2, 3+4 or 1+4, Figure 2) as outlined above. S2R+ attP 99F8 Clone 141 was further expanded and RMCE experiments were carried out. This clone is available from DGRC (DGRC #315). Single cell clones were derived using the two-step sorting protocol (Bulk sorting followed by single cell sorting) detailed above for the following lines: S2-DGRC-attP-25C6-Clone 8 (DGRC # 316), Kc167-attP-25C6-Clone 35 (DGRC Cat# 317), Kc167-attP-99F8-Clone 71 (DGRC Cat# 318), ML-DmBG2-c2-attP-25C6-Clone 8 (DGRC # 319). Single cell clones for ML-DmBG3-c2-attP-25C6-Clone 74 (DGRC Cat# 320) and ML-DmBG3-c2-attP-99F8-Clone 28 (DGRC # 321) were derived directly by one-step single cell sorting.

### Immunostaining

To confirm dsRed/EGFP expression, cells were plated onto poly-lysine coated chamber slides (Thermoscientific Nunc Lab-Tek), allowed to attach for 1-2 hours, fixed with 4% paraformaldehyde in PBS, pH 7.0, washed and immunostained with rabbit Anti-dsRed (632496, Clontech; 1:1000) and/or mouse Anti-EGFP (DHSB 12A6) antibodies and the respective Alexa dye-coupled secondary antibodies (Jackson ImmunoResearch) reconstituted in 5% Normal Goat Serum, 0.1% Triton in PBS. Cells were then mounted on DAPI containing mounting media (Vectashield H-1200) prior to imaging on a Leica SP8 confocal microscope.

### Data availability

All data necessary for confirming the conclusions in this paper are included in this article and in supplemental figures and tables. All cell lines and plasmids are available upon request from the DGRC.

## Results and Discussion

### Generation of single cell clones with 99F8 attP Act::dsRed insertion

To determine the optimal loci to introduce a docking cassette with flanking attP sites in the 3^rd^ chromosome, we first explored which of the attP *Drosophila melanogaster* lines had the highest survival rates of transformants listed in the Rainbow Transgenics website. Amongst the lines that were positive for transgenic transformants, the insertion at 99F8 on 3R had the highest survival rate (Figure 1A). Subsequently, the 99F8 locus around the *in vivo* attP insertion was sequenced in S2R+, Kc167, ML-DmD17 and ML-DmBG3-c2 cell lines and examined for variability and consensus between cell lines. Apart from a single nucleotide polymorphism in S2R+, the sequence around the intended attP insertion site was conserved across all the cell lines compared to the *Drosophila melanogaster* genomic sequence (Flybase release 6.32, Figures 1B and 1C). We therefore chose the 99F8 locus on 3R as the target for attP site insertion.

**Figure 1.**
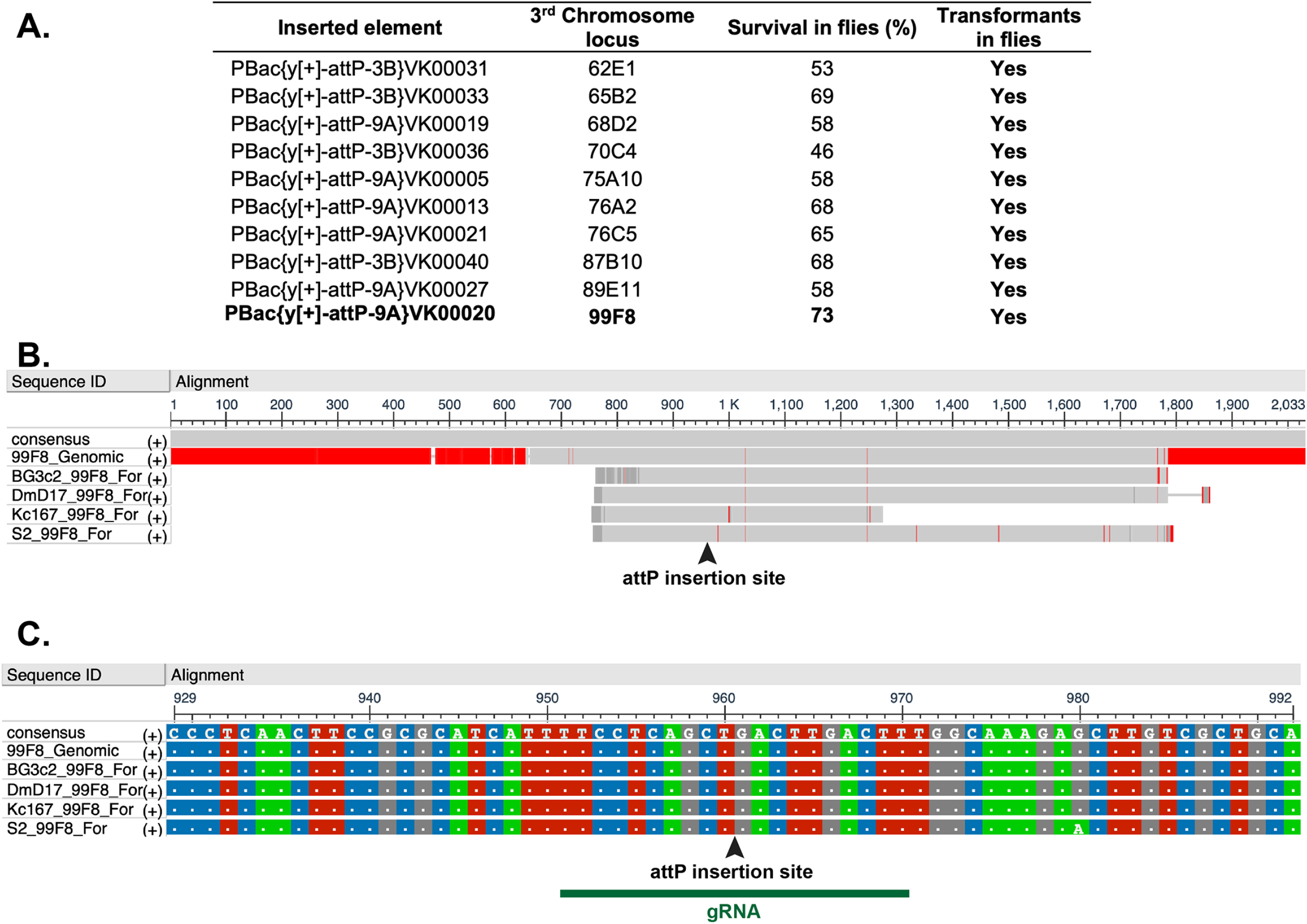
Third chromosome locus 99F8 is ideal for attP site insertion. (**A**) Comparison of attP fly lines available from Rainbow Transgenics Inc. outlining the chromosomal position and percent viability. Only the lines positive for producing transformants have been listed. The locus 99F8 chosen for generation of attP lines is shown in bold (**B**) Multiple sequence alignment of the 99F8 genomic locus (Flybase release 6.32) with sequencing data from genomic DNA PCRs amplifying the 99F8 locus in ML-DmBG3-c2, ML-DmD17, Kc167 and S2R+ cell lines, red indicates absence of consensus. (**C**) A zoom in view of the multiple sequence alignment from **B** demonstrates overall consensus except for a single nucleotide in S2R+ (at position 960). The arrowhead in (B) and (C) specifies the insertion site with the green bar outlining the gRNA target site (Table 1).

To generate the attP insertion at the 99F8 locus using CRISPR-Cas9 strategy, cells were transfected with a single gRNA (Figures 1B and 1C) accompanied by the repair construct containing the attP>>Act::dsRed<<attP cassette flanked with ∼1 kb homology arms. Cells positive for dsRed were sorted by fluorescence activated cell sorting (FACS) 19 days after the CRISPR-Cas9 transfection. A two-step FACS approach was adopted to derive single cell clones. In the first step, a bulk sort of dsRed positive cells from the CRISPR-Cas9 transfection sorted the cells into a single well of a 6-well plate into S2R+ conditioned medium. A total of 4,422 dsRed positive cells, 0.3% of the sorted population were pooled together. Following expansion of the dsRed positive pooled cells for 20 days, potential integration at 99F8 locus was ascertained by PCR amplification from a sample of the positive dsRed pooled cells and sequence confirmation of the insert junctions (Figure S1A). For the second single cell cloning FACS step, the amplified bulk pooled cells were subjected to FACS to sort single dsRed positive cells into each well of a 96-well plate. In contrast to the bulk sorting step, 64.7% of the sorted cells were dsRed positive. Single cell colonies were observed in 18% (258/1440) of the wells, of which 60% (154/258) of the clones were positive for dsRed expression (Table 2). Integration of the attP>>Act::dsRed<<attP cassette was assessed by PCR with primers that amplified across the integration locus from outside the homology arms for each single cell colony (Figure 2A, primers 1 and 4). A successful insertion yielded a 5.1 kb amplicon whereas the presence of a 1.5 kb amplicon indicated that the 99F8 genomic locus remained unmodified (Figure S1B). Of the 151 dsRed positive single cell clones assessed, 26 were positive for the 5.1 kb amplicon. However, all of the clones positive for the 5.1 kb amplicon also possessed the 1.5 kb amplicon indicating that that there was at least one unmodified 99F8 locus in these lines (Figure S1B). There are at least four copies of the 3^rd^ chromosome in S2R+ cells (Lee *et al*. 2014; Senaratne *et al*. 2016), and as a result, all of these clones could have anywhere between one to three modified 99F8 loci. Further PCR typing revealed that only one out of the 26 clones, clone 141, tested positive for all the expected combinations of PCR amplicons wherein primers from within the inserted cassette and outside the homology arms were used (Figure 2B). The authenticities of the amplicons were confirmed by sequencing and thus the S2R+ Clone 141 was validated to contain the attP>>Act::dsRed<<attP cassette. The S2R+ Clone 141 is therefore heterozygous for the 99F8 attP site.

**Figure 2.**
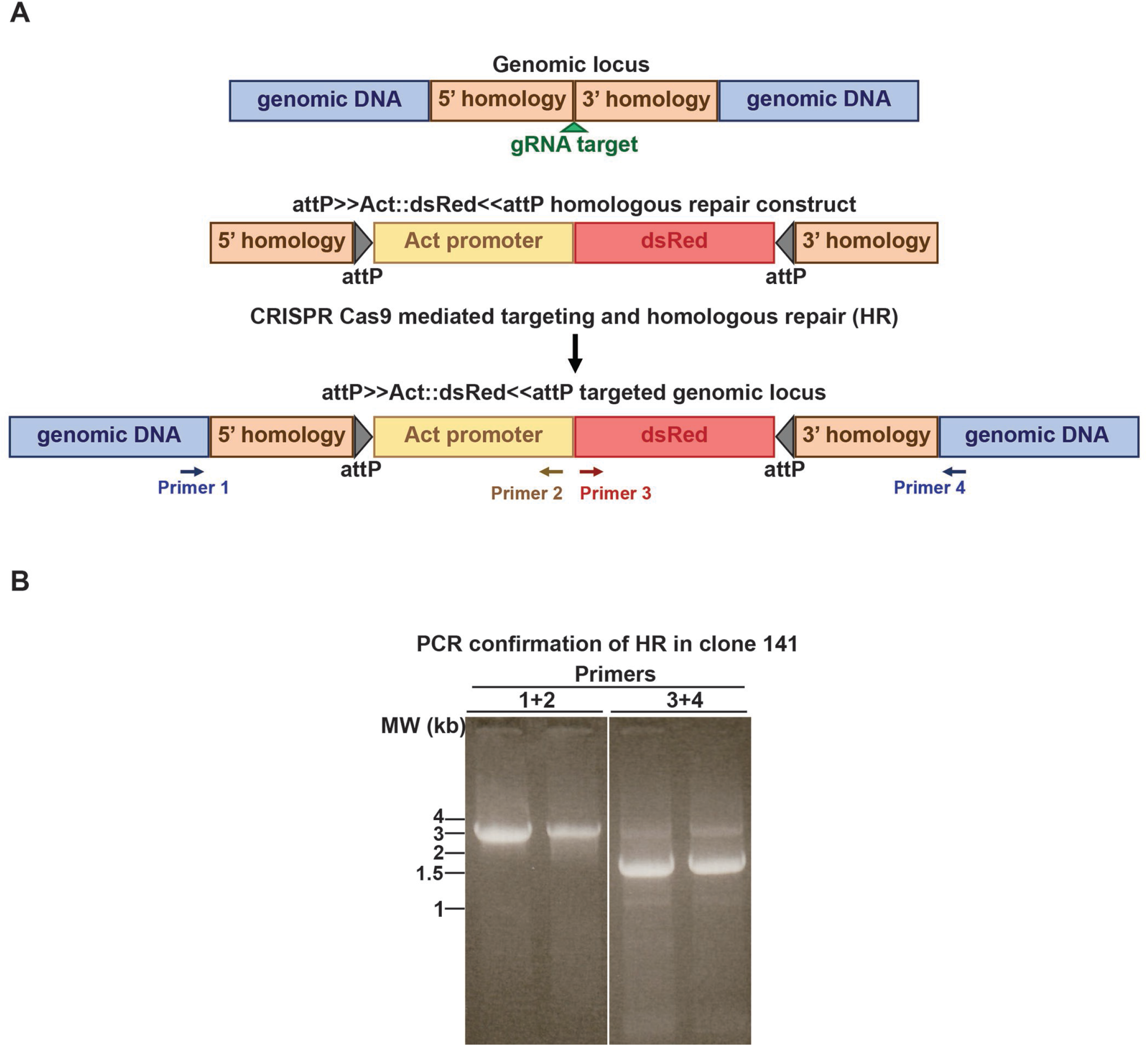
Generation of 99F8 attP site in S2R+ cell line using a CRISPR-Cas9 strategy. **(A)** Approximate 1 kb long 5’ and 3’ homology arms flanking the gRNA target/insertion site was used to create the attP>>Act::dsRed<<attP homologous repair (HR) cassette. Upon successful homologous repair the 99F8 locus is expected to contain the attP sites and the cells will ubiquitously express dsRed under the control of the Actin promoter. **(B)** Successful HR was confirmed by PCR utilizing the primer pairs 1 and 2 or 3 and 4 (see **A**) that generated amplicons of 3.3 and 1.7 kb respectively, as seen in the S2R+-attP-99F8-Clone 141 line. Shown are duplicate samples that from the same reaction. All the amplicons obtained were sequence verified.

**Table 2.** Process efficiency in obtaining single cell clones containing the expected insertion at the 25C6 locus for distinct *Drosophila* cell lines.

| Cell Line | Genomic locus | Single cell clones | PCR analysis: Putative (+) clones | Sequence verified | Heterozygote/ Homozygote |
| --- | --- | --- | --- | --- | --- |
| S2-DGRC | 25C6 | 9 / 960 (1 %) | 1 /9 | 1 | Homozygote |
| Kc167 | 25C6 | 447/960 (47%) | 9/200 | 1 | Homozygote |
| ML-DmBG2-c2 | 25C6 | 13/ 960 (1.35%) | 5/13 | 1 | Homozygote |
| ML-DmBG3-c2 | 25C6 | 83 / 960 (8.6 %) | 19/ 83 | 3 | Homozygote:1<br>Heterozygote: 2 |
| S2R+ | 99F8 | 258/1440 (18 %) | 8/153 | 1 | Heterozygote |
| Kc167 | 99F8 | 255/1056 (24%) | 7/204 | 1 | Heterozygote |
| ML-DmBG3-c2 | 99F8 | 136/1056 (13 %) | 7/38 | 2 | Heterozygotes |

The two-step FACS procedure (Cherbas *et al*. 2015) significantly improved the fraction of dsRed positive cells available for the cloning step. Moreover, the extended time from transfection to cloning meant reduced numbers of dsRed positive transiently transfected cells. Nevertheless, the overall time to generate these cell lines, from CRISPR-Cas9 transfection to freezing aliquots of confirmed single cell clones was just over 4 months. This can be accelerated in instances where documenting and clarifying the status of each of the single cell clone derived is not essential. Use of drug selection in the previous approaches (Cherbas *et al*. 2015; Manivannan *et al*. 2015) inadvertently also selects for insertion at non-target loci. With the use of minimal plasmid amounts during transfection and elimination of drug-based selection in the current approach, integrations that may occur at non-target loci are mitigated and subsequently eliminated by single cell cloning.

### Recombination mediated cassette exchange demonstrates the functionality of the S2R+-attP-99F8 line

Using the S2R+-attP-99F8 Clone 141 line, RMCE was attempted with an attB>>Act::GFP<<attB cassette (Figure 3A). Following RMCE, we expected to observe three population of cells: dsRed positive (cells without successful RMCE), GFP positive (cells with all the copies of Act::dsRed replaced by RMCE with Act::GFP) and double positive cells (cells with successful RMCE in at least one of the third chromosomes at one 99F8 locus, but retaining the Act::dsRed in at least one other third chromosome, Figure 3B). Among the FACS sorted cell population, 1.7% cells were GFP positive, but negative for dsRed. Double positive cells accounted for 7.5% of the sorted population (Figure 3B). The design of the attP sites in the S2R+ attP line permits for cassette exchange in either orientation. Genomic DNA from the dsRed positive, GFP positive and the double positive fractions was used to confirm the RMCE. Cassette exchange was confirmed with PCR wherein one primer from within GFP was used in combination with either the upstream (Primer 7, Figure 3A) or downstream (Primer 6, Figure 3A) primers. The dsRed positive fraction was negative for both sets of PCRs, while amplicons of the expected size were observed for GFP positive and double positive fractions (Figure 3C). Sequencing the amplicons confirmed their authenticity. This demonstrated that bidirectional cassette exchange can be achieved using the S2R+-99F8-attP line. To obtain a clonal population, it would however be imperative to isolate single cell clones by FACS. There is also the possibility that not all of the attP sites in S2R+-attP-99F8 line has had a successful RMCE, hence the presence of dsRed and GFP double positive cells harboring both the Act::dsRed and Act::GFP cassettes. Additionally, given the data, it is possible that two attP loci with a single cell have the inserted Act::GFP cassette in both orientations. Alternatively, the RMCE confirmation results (Figure 3C) demonstrating bidirectional Act::GFP cassette exchange could be the result of separate population of cells carrying the cassette oriented in either direction.

**Figure 3.**
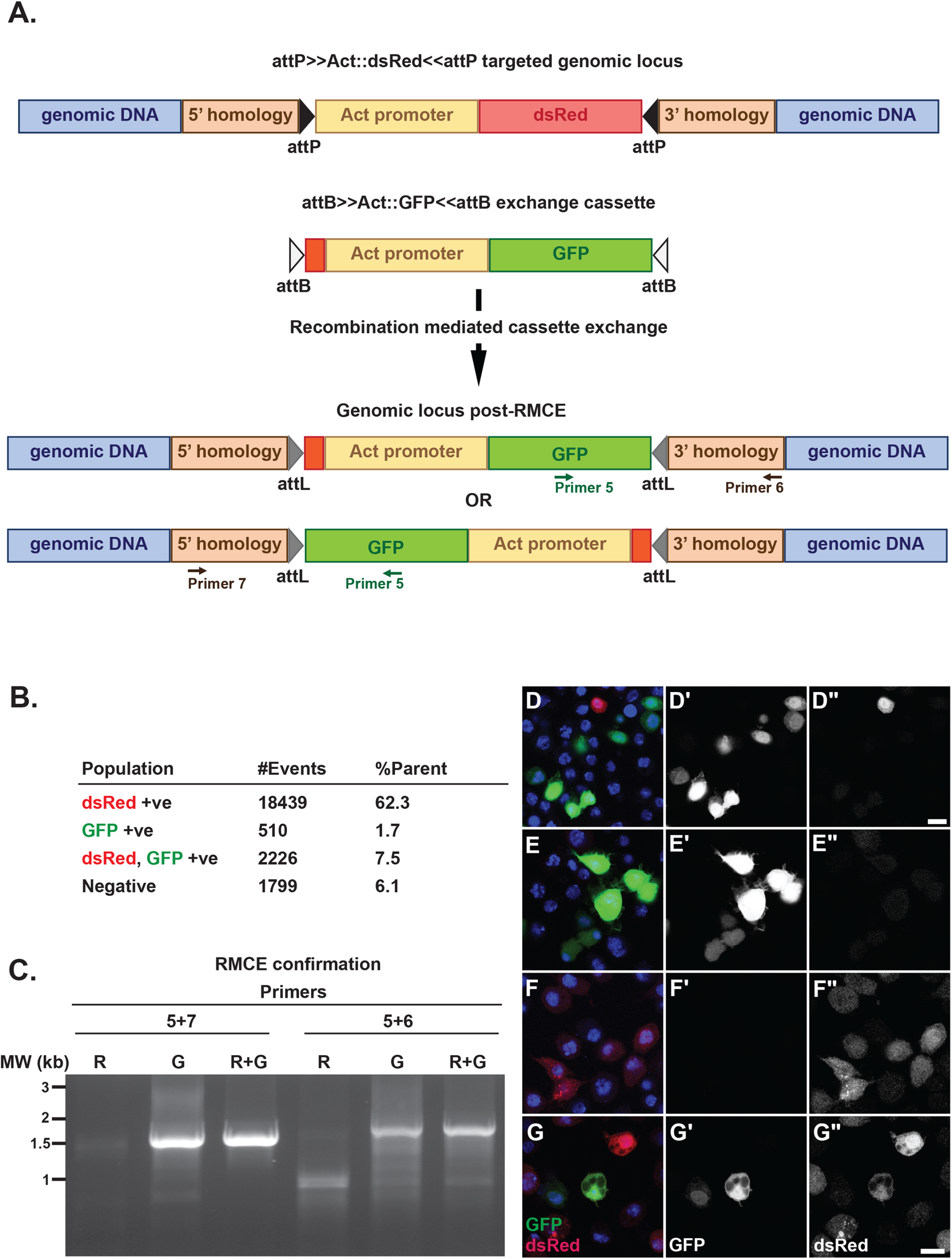
Confirmation of successful cassette exchange at the 99F8 attP site in S2R+-attP-99F8 line. **(A)** A recombination mediated cassette exchange (RMCE) construct with attB>>Act::GFP<<attB was transfected with PhiC31 integrase to generate cells that exchanged the dsRed for GFP at the 99F8 attP locus. The exchanged cassette can insert in either orientation. **(B)** Bulk sorting the RMCE cassette transfected cells by FACS revealed that 1.7% of the cells were positive for GFP alone indicating that all the copies of Act::dsRed were exchanged with Act::GFP in this fraction of cells. #Events is the number of dsRed only, GFP only or dsRed, GFP double positive cells detected by FACS with the respective percentages indicated in the %Parent column. **(C)** Successful RMCE in either orientation was diagnosed using the primer pairs 5+7 or 5+6 (see **A**) that generated amplicons of 1.5 and 1.7 kb, respectively. The following fractions obtained from FACS were assessed for RMCE: dsRed positive (R, cells without successful RMCE), GFP positive (G, cells with all the copies of Act::dsRed replaced by RMCE with Act::GFP) and double positive (RG, cells with successful RMCE in at least one of the chromosomes at the 99F8 locus, but retaining the Act::dsRed in at least one other homolog). (**D**) Transfected cells before FACS, (**E**) GFP positive, (**F**) dsRed positive and (**G**) double positive cell fractions after FACS sorting were immunostained with anti-GFP (D′, E′, F′, G′) or anti-dsRed (D′′, E′′, F′′, G′′) antibodies. Nuclei are stained with DAPI (blue). The respective merged images are shown in D, E, F and G. Scale bar = 10 μm

Investigating the expression of GFP and dsRed by immunostaining revealed that the cells in GFP positive (Figure 3E,E′,E′′) and dsRed positive (Figure 3F,F′,F′′) fractions only expressed either GFP or dsRed, respectively. In the double positive fraction, cell populations exclusively expressing either GFP or dsRed and those expressing both GFP and dsRed were observed (Figure 3G, G′, G′′), suggesting sorting was not absolute. Given that the S2R+-attP-99F8 clone 141 has at least one unmodified 99F8 locus, the immunostaining data indicates that at least two of the four 3^rd^ chromosomes in the S2R+-attP-99F8 line carry the attP site. As there is one 99F8 locus that is unmodified (Figure S1B) in the S2R+-attP-99F8 line, the detection of both the GFP and dsRed signals in the same cell indicates a minimum of two 99F8 attP sites.

Constitutive overexpression of certain transgenes in cell culture could potentially be toxic leading to cell death. This can be circumvented by the use of an inducible metallothionein (MT) promoter (Bunch *et al*. 1988; Klueg *et al*. 2002). We therefore tested another exchange cassette with the MT promoter, attB>>MT::GFP<<attB for RMCE in the S2R+-attP-99F8 cell line. In contrast to transfection with the attB>>Act::GFP<<attB construct that constitutively expressed GFP, GFP expression in attB>>MT::GFP<<attB is induced by the presence of Cu^2+^. Consequently, the GFP- and double-positive cells upon FACS sorting the Cu^2+^ induced, transfected cells accounted for only 0.3% and 1.7% of the sorted population (Figure S2A). Despite the low percentages of the GFP- and double positive fractions, it was feasible to expand these fractions to an extent that cassette exchange at the appropriate 99F8 locus (Figure S2B) could be confirmed and sequence verified.

It is evident from the data presented that the inserted attP site(s) in the S2R+-attP-99F8 Clone 141 line are functional. Taken together, it is most likely that this line has at least 2-3 attP insertions at the 99F8 locus given the number of copies of the 3^rd^ chromosome in S2R+ cells (Lee *et al*. 2014; Senaratne *et al*. 2016). Potentially, this feature allows for multiple rounds of RMCE with unique constructs, thereby creating double or possibly triple transgenics in the same cell line with rest of the genome unchanged. Alternatively, it is likely that in some clones all the attP sites have undergone RMCE, thereby yielding lines with more robust transgene expression. These results suggest the possibility that a series of clones with increasing transgenic copy numbers could be derived, which would be useful in experiments requiring a battery of varying transgene expression.

### Generation of multiple attP lines

To expand the choice of attP sites, another locus at 25C6 on the second chromosome was chosen to introduce an attP site in S2-DGRC cell line. Like the 99F8 site, we determined the genomic sequence at 25C6 is conserved across five different cell lines from three different genetic backgrounds: S2-DGRC, Kc167, and the Miyake laboratory CNS-derived lines: ML-DmD17-c3, ML-DmBG2-c2 and ML-DmBG3-c2 (Figure S3). To generate the attP insertion at the 25C6 locus using CRISPR-Cas9 strategy, cells were transfected with two gRNAs accompanied by the repair construct containing the attP>>Act::dsRed<<attP cassette flanked with the homology arms. Cells positive for dsRed were enriched by fluorescence cell sorting at least 10-20 days after the initial transfection before sorting to derive single cell clones. The integration of the donor repair cassette was assessed by PCR amplification across the integration locus from outside the homology arms (Figure S4A). A successful insertion yielded a 5.7 kb amplicon while a 2.6 kb amplicon was observed if the 25C6 genomic locus remained unmodified. A non-homozygous modification of the locus would be expected to yield both bands, while a homozygous genomic modification would only yield a single 5.7 kb amplicon (Band 1) (Figure S4A). We further used primer pairs that amplified from within insert to outside the homology arms (Band 2, 4.7 kb and Band 3, 4.5 kb) to verify the clones and also sequence for the presence of the attP sites (Figure S4A). This PCR diagnosis indicated that we obtained a homozygous S2-DGRC clone containing the attP cassette at the 25C6 locus (Figure S4B).

In order to investigate the applicability of this technique to other *D. melanogaster* cell lines, we introduced attP sites at 25C6 and 99F8 in other cell lines. We documented the homozygote Dm-MLBG2-c2-attP-25C6-Clone 8 (Figure S4C) and ML-DmBG3-c2-25C6-Clone 74 (Figure S4D), Kc167-attP-25C6-Clone 35 (Figure S5A) and the heterozygote Kc167-attP-99F8-Clone 71 (Figure S5C) and ML-DmBG3-c2-attP-99F8 28(Figure S6). We summarized the various efficiencies of the genomic modification process and single cell cloning with different cell lines in Table 2. The current status of the multiple attP cell lines that have been generated at DGRC using this CRISPR-Cas9 approach are shown in Table 3. In addition to generating the attP lines at both the chromosomal loci in Kc167 cells (Figure S5D,E), we have currently demonstrated successful RMCE utilizing the attB>>Act::GFP<<attB cassette in Kc167 at both 25C6 and 99F8 loci (Figure S5C, S5D). As noted, the inverted attP sequences flanking the docking site allows the bi-directional insertion of the donor cassette. After RMCE, we detected in the enriched population of GFP positive cells in which the cassette was inserted in both orientations (Figure S5C, S5D).

**Table 3.**
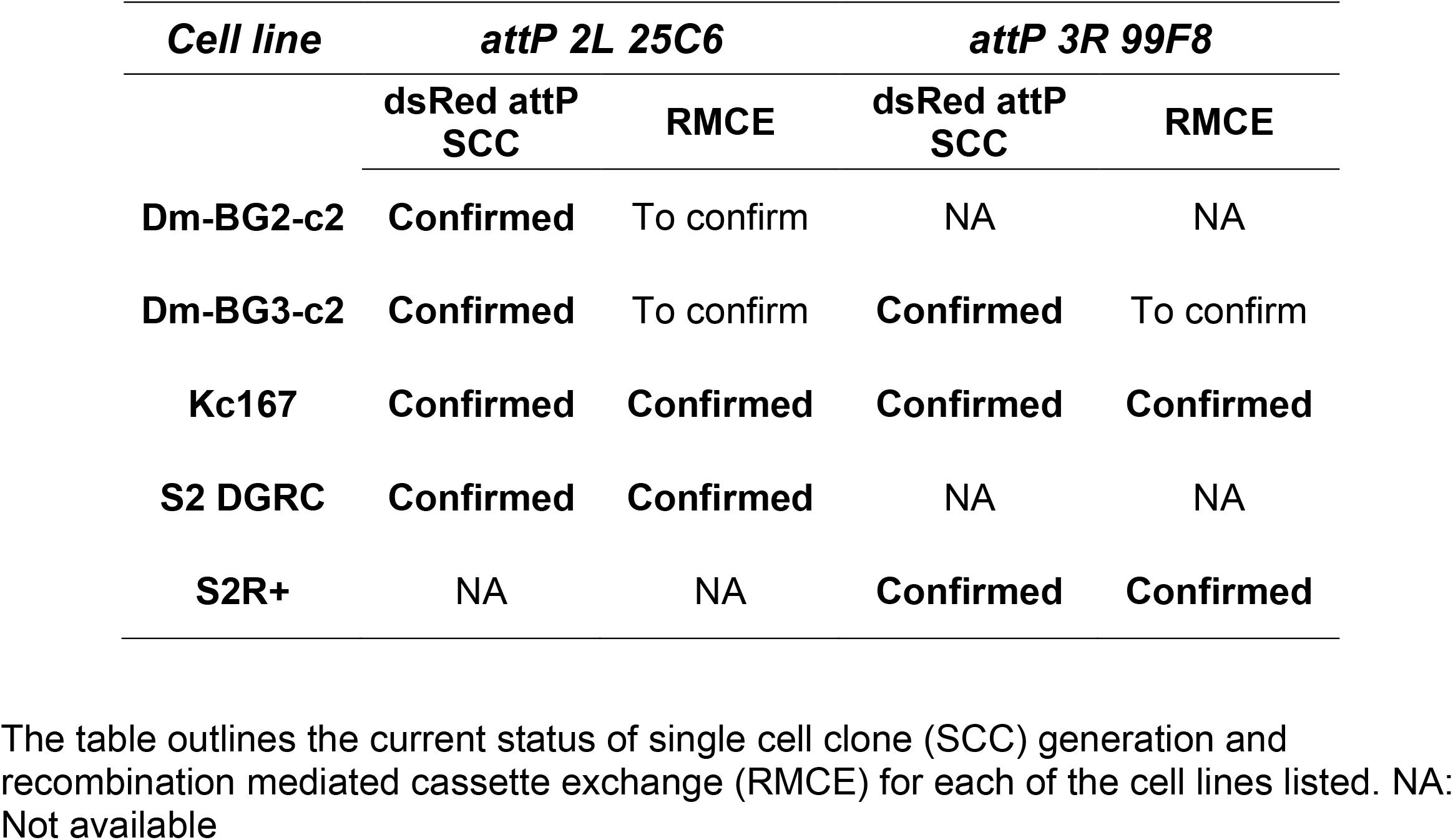
Summary of the various attP>>Act5C::dsRed<<attP cell lines generated at 25C6 and 99F8 genomic loci.

In summary, we have been able to insert RMCE docking cassettes containing flanking inverted attP sites in two different genomic loci in both embryonic and larval CNS derived cell lines. Successful cassette exchange was demonstrated in the generated S2 and Kc167 attP lines. Moreover, the methodology presented here not only permits the introduction of attP sites at loci 25C6 and 99F8 in any cell line but also with a few minor modifications of the reagents generated in this study, an attP site could be inserted in any desired locus. Interestingly, our results demonstrate that the insertion of an attP site in every copy of the genome and RMCE conversion is variable. Nonetheless, this “heterozygosity” would enable researchers to generate a multitude of clones that could have multiple transgenes within the same clonal line or express varying doses of the transgene with the option to choose the directionality of the exchanged cassette. In addition, our approach now permits the comparisons of transgenic expression from the same locus across various biological and physiologically diverse *D. melanogaster* cell lines. All the reagents generated in this study will be accessible to researchers from the *Drosophila* Genomics Resource Center to create their own cell lines, as well as a cost-recovery service to aid with deriving clonal populations post-RMCE. Overall, these cell lines and plasmid collection created will aid researchers to further harness *D. melanogaster* cell culture to address broader research questions.

## Acknowledgments

We thank Dr. Kris Klueg for critical input in shaping the manuscript. We thank the Light Microscopy Imaging Center (Indiana University), Gregory P. Crouch and Amanda LJ. McKeen for help with irradiating cells, Christiane Hassel and the IUB Flow Cytometry Core Facility (Indiana University). This work is supported by a NIH grant (2P40OD010949) to the *Drosophila* Genomics Resource Center.

---

**Figure S1.**
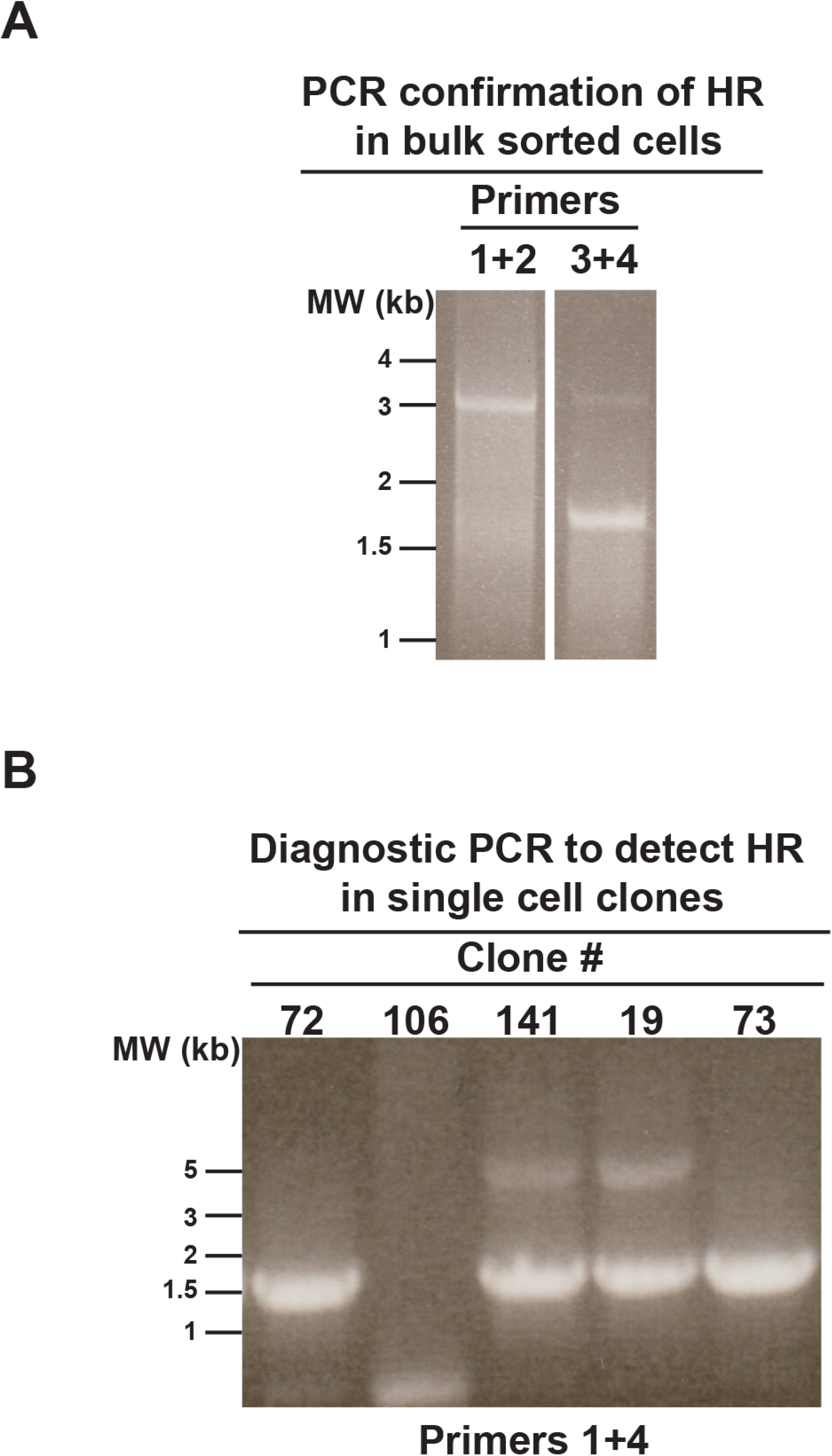
PCR diagnosis of CRISPR homologous recombination in S2R+ cells. (A) Homologous recombination after CRISPR was diagnosed with primer pairs 1+2 and 3+4 (Figure 2). (**B**) Initial screening for all the putative single cell clones was performed with the primer pair 1+4 (Figure 2). Clones positive (91, 141) for successful HR yielded a 5.1 kb amplicon in addition to the unmodified 1.5 kb amplicon, indicating insertion of the attP>>Act::GFP<<attP cassette.

**Figure S2.**
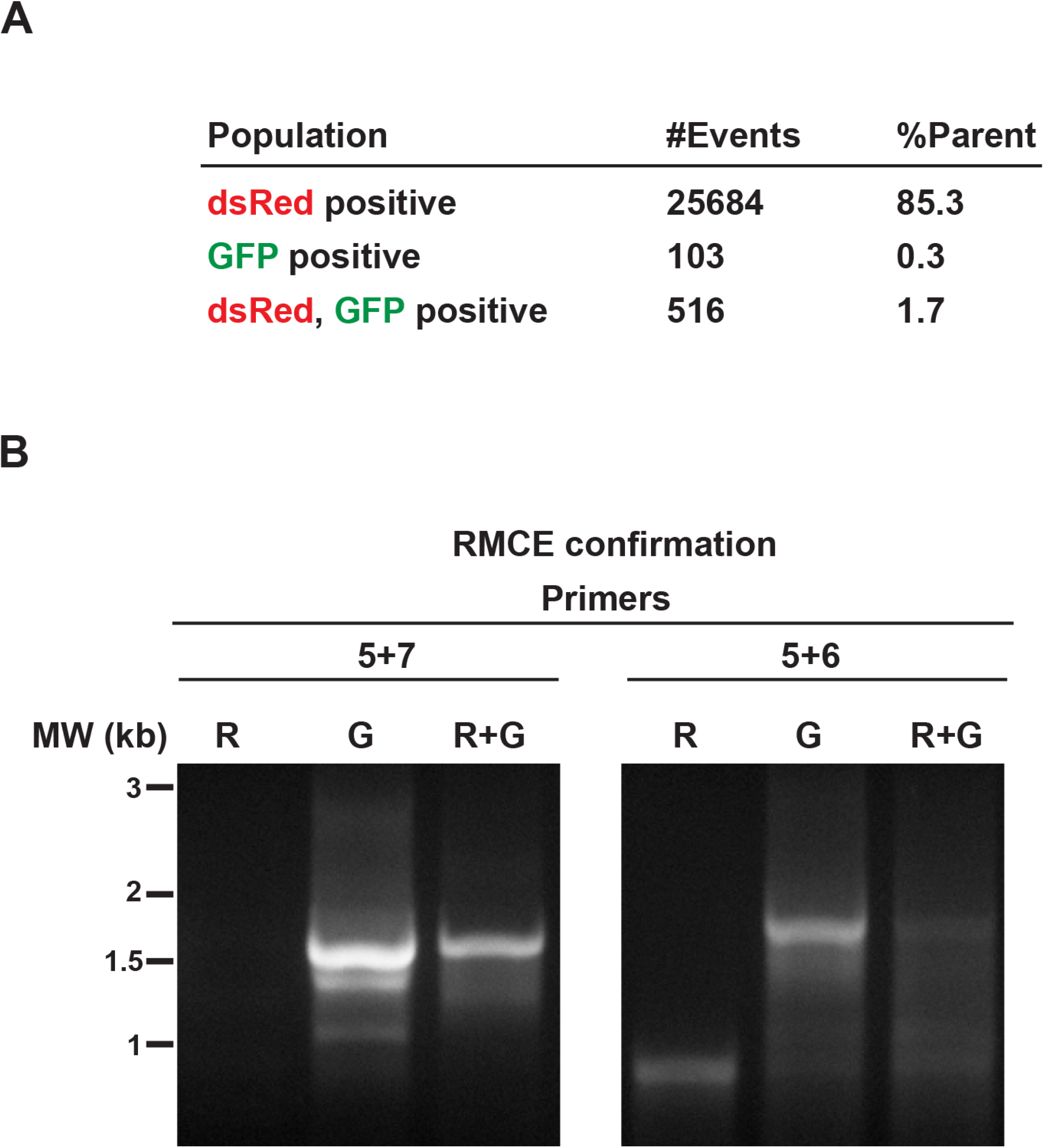
Recombination mediated cassette exchange using MT::GFP cassette in S2R+-attP-99F8-Clone 141. **(A)** Bulk sorting the S2R+ 99F8 attP cells after transfecting with attB>>MT::GFP<<attB recombination mediated cassette exchange (RMCE) cassette by FACS revealed that 0.3% of the cells were positive for GFP alone indicating that all the copies of Act::dsRed were exchanged with MT::GFP in this fraction of cells. #Events is the number of dsRed only, GFP only or dsRed, GFP double positive cells detected by FACS with the respective percentage indicated in the %Parent column. **(B)** Successful RMCE in either orientation was diagnosed using the primer pairs 5+7 or 5+6 (Figure 3) that generated amplicons of 1.5 and 1.7 kb, respectively. The following fractions obtained from FACS were assessed for RMCE: dsRed positive (R, cells without successful RMCE), GFP positive (G, cells with all the copies of Act::dsRed replaced by RMCE with MT::GFP) and double positive (RG, cells with successful RMCE in at least one of the chromosomes at the 99F8 locus, but retaining the Act::dsRed in at least one other homolg).

**Figure S3.**
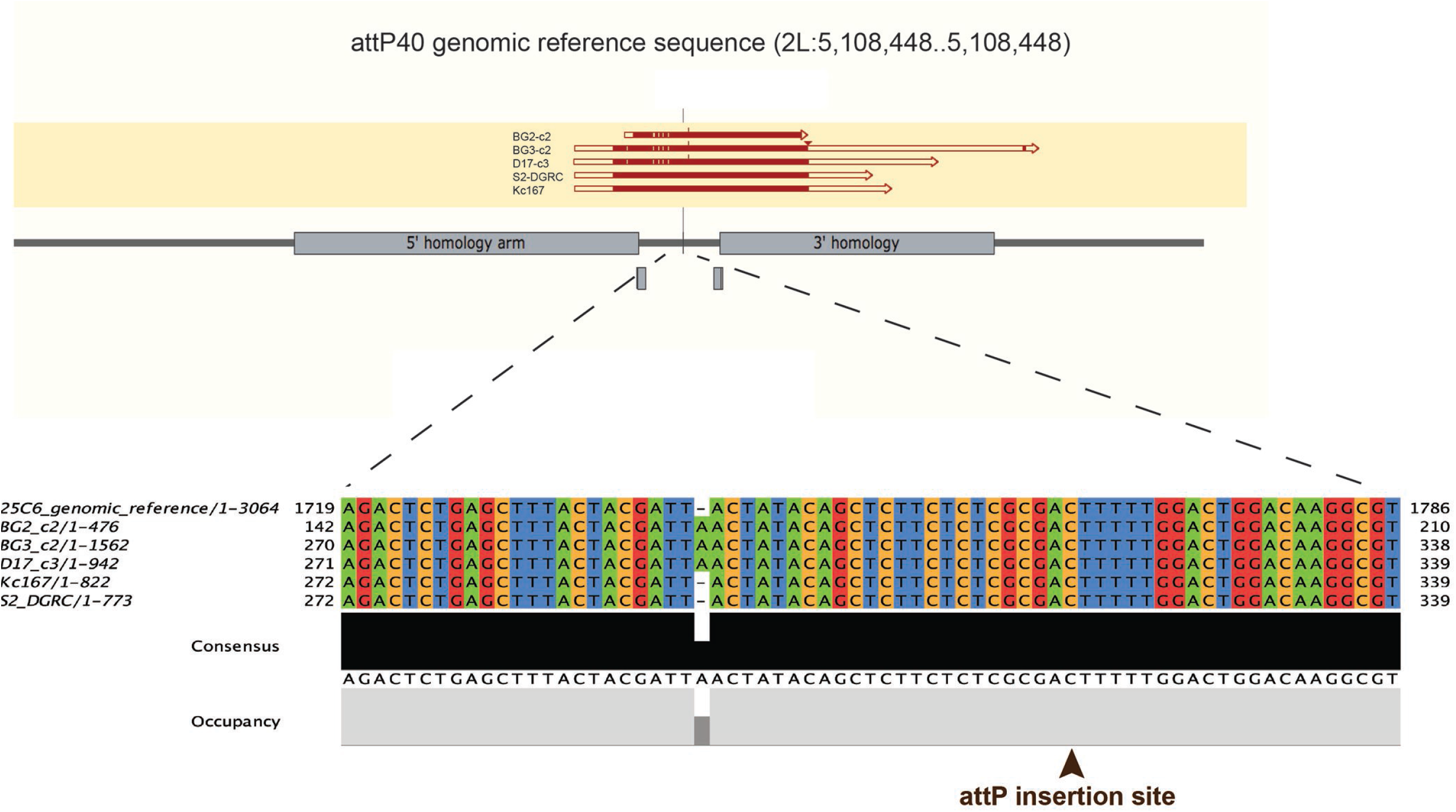
Sequence alignment of the genomic DNA from multiple Drosophila melanogaster cell lines derived from different genetic backgrounds at the 25C6 locus(2L:5,108,448..5,108,448). Multiple sequence alignment of the 25C6 genomic locus for the following lines: ML-DmBG2-c2, ML-DmBG3-c2, ML-DmD17-c3, Kc167 and S2-DGRC. The sequences were conserved at the two gRNA cut sites. A zoomed in view of the multiple sequence alignment demonstrated the overall consensus.

**Figure S4.**
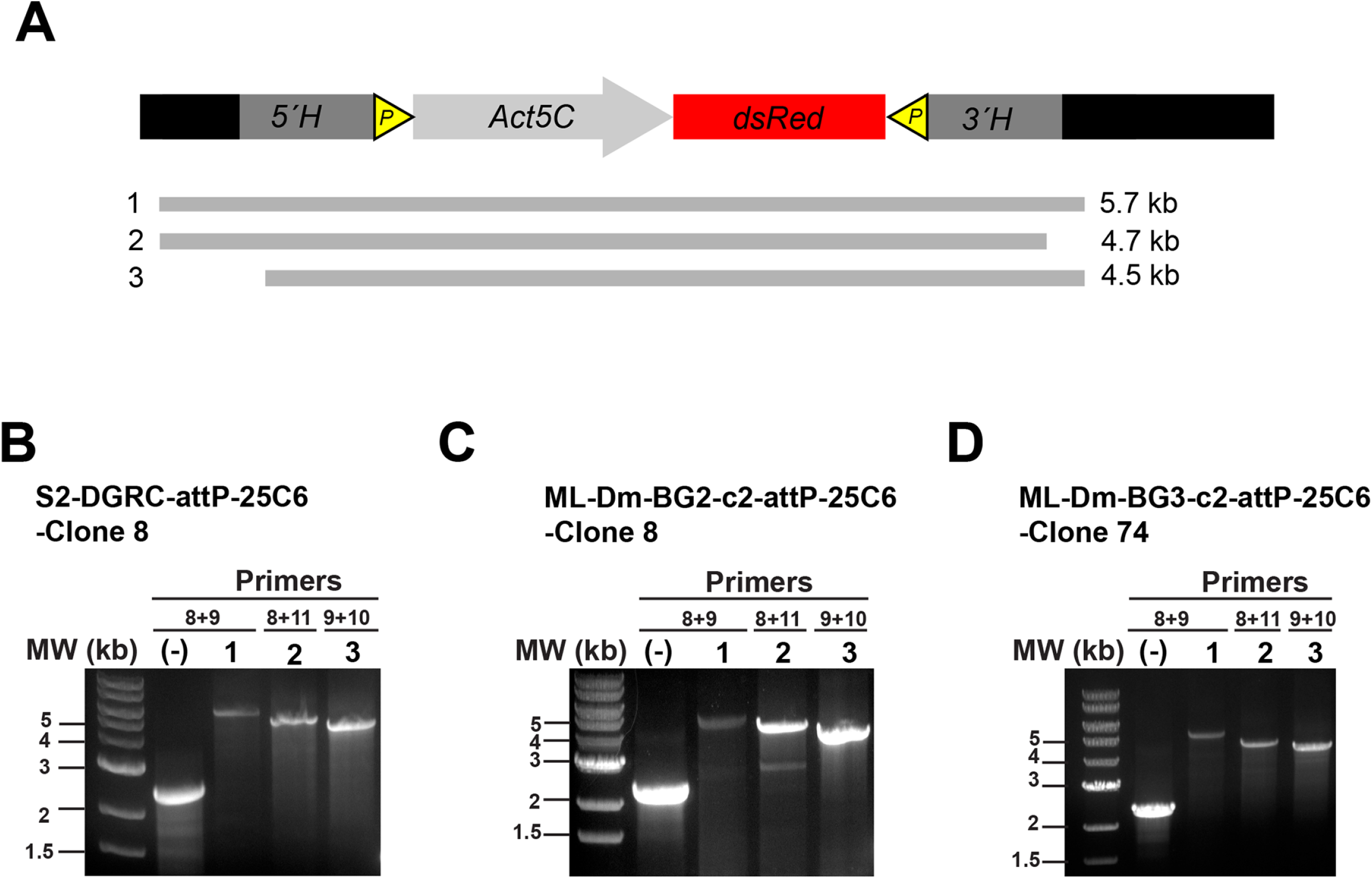
Generation of 25C6 attP site in *D. melanogaster* cell lines using CRISPR-Cas9. (**A)** A schematic of the insertion at the 25C6 locus, indicating the regions outside the insert homology arms (black), 5’ homology arm of the insert (5’H), attP site (yellow triangle), act5C promoter (gray arrow), dsRed (red bar) and the 3’ homology arm of the insert (3’H). The numbered gray bars (1-3) correspond to the sizes of the PCR products for verifying the insertion. Bar 1 was amplified using Primer 8 + 9. Ba r 2 is amplified with Primers 8 + 11. Bar 3 was amplified using primer 9 + 10. (**B**) PCR verification of S2-DGRC-attP-25C6-Clone 8, (-) control is the parental S2-DGRC line amplified with the primer pairs outside the homology arms. (**C**) PCR verification of ML-Dm-BG2-c2-attP-25C6-Clone 8, (-) control is the parental ML-Dm-BG2-c2 line amplified with the primer pairs outside the homology arms. (**D**) PCR verification of ML-Dm-BG3-c2 attP-25C6-Clone 74, (-) control is the parental ML-Dm-BG3-c2 line amplified with the primer pairs outside the homology arms.

**Figure S5.**
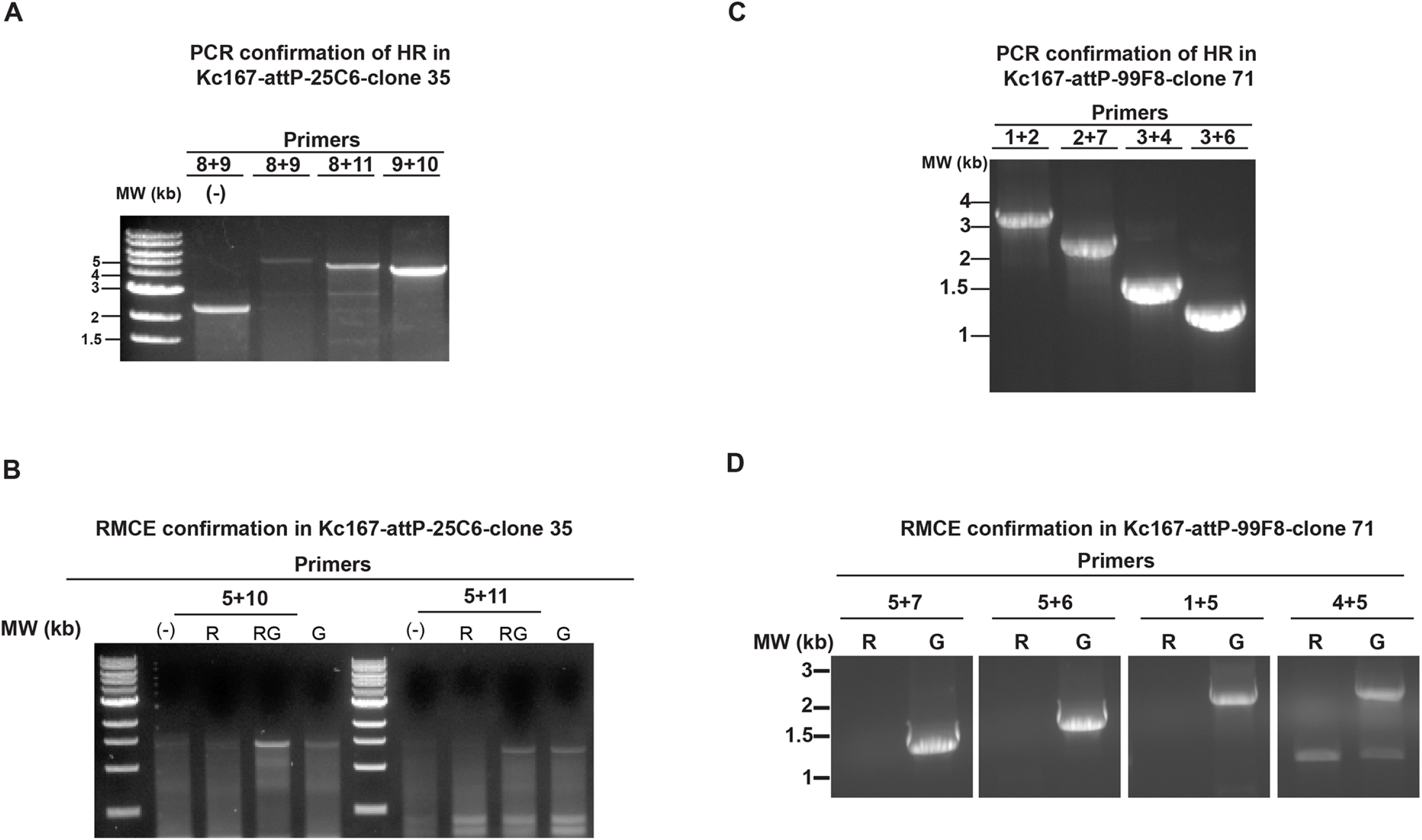
Generation of Kc167-attP-25C6 and Kc167-attP-99F8 lines and RMCE confirmation. (**A**) Homologous recombination of the cassette after CRISPR was detected at 25C6 locus with primer pairs 8+9, 8+11 and 9+10 in clone 35 (Table 1). (-) control is the parental Kc167 line (**B**) Successful RMCE was diagnosed using either the primer pair 5+10 or 5+11, which indicated that the donor cassette was inserted in reverse (3’ to 5’) and forward (5’ to 3’) orientations, respectively. (-) control is the parental Kc167 amplified by primer pair 5+10 and primer pair 5+11, respectively. The following fractions obtained from FACS were assessed for RMCE: dsRed positive (R, cells without successful RMCE), GFP positive (G, cells with all the copies of Act::dsRed replaced by RMCE with Act::GFP) and double positive (RG, cells with successful RMCE in at least one of the chromosomes at the 25C6 locus, but retaining the Act::dsRed in the other homolog). (**C**) Homologous recombination after CRISPR was diagnosed at 99F8 locus with primer pairs 1+2, 2+7, 3+4 and 3+6 (Figures 2 and 3) in clone 71. (**D**) Successful RMCE was diagnosed using the primer pairs 5+7, 5+6, 1+5 or 4+5 (Figures 2 and 3), dsRed positive (R, cells without successful RMCE) and GFP positive (G, cells with all the copies of Act::dsRed replaced by RMCE with Act::GFP).

**Figure S6.**
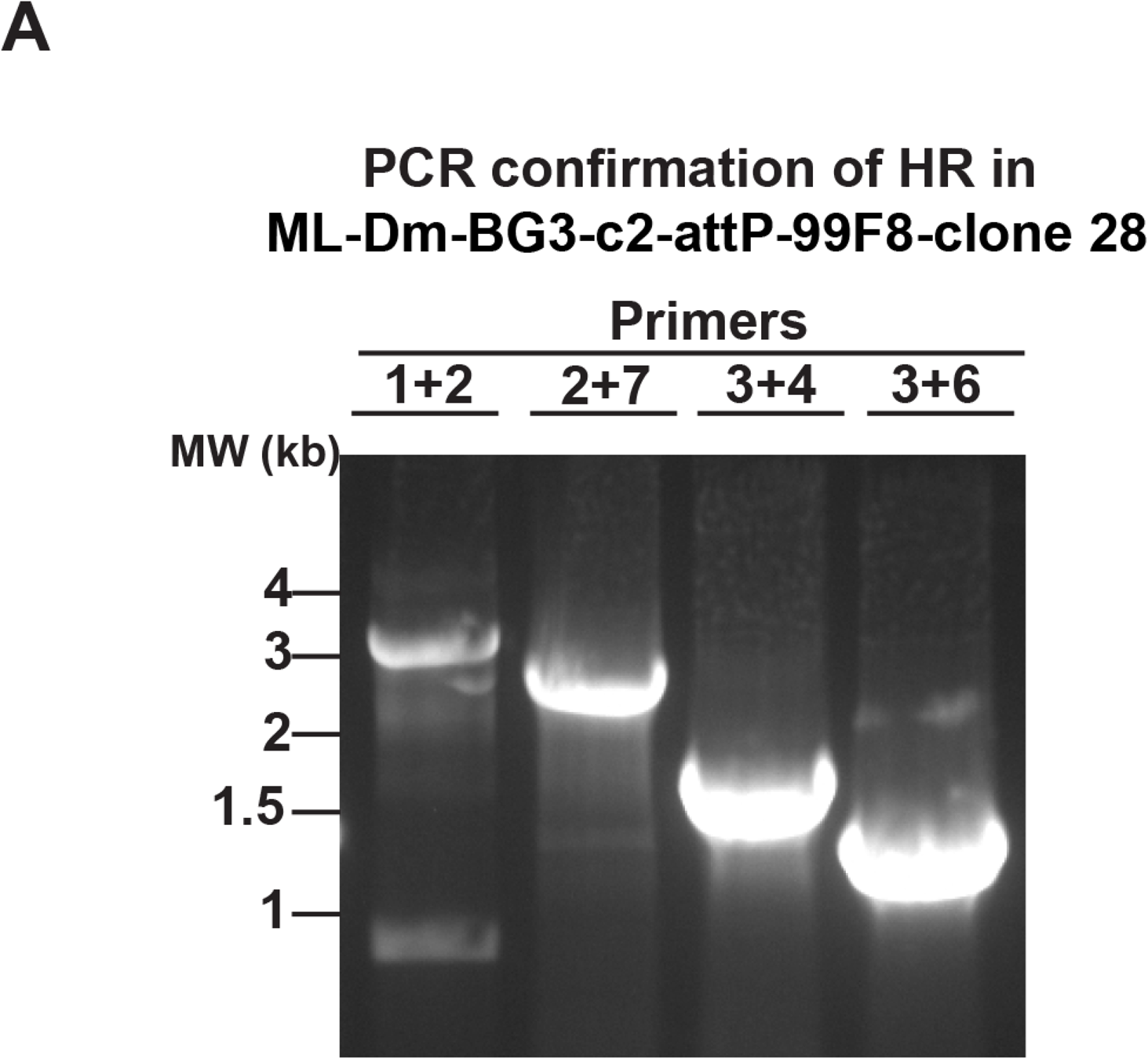
Generation of ML-Dm-BG3-c2 -attP-99F8 line. Homologous recombination after CRISPR editing was diagnosed at 99F8 locus with primer pairs 1+2, 2+7, 3+4 and 3+6 (Figures 2 and 3) in clone 28.

